# DISCOVER-EEG: an open, fully automated EEG pipeline for biomarker discovery in clinical neuroscience

**DOI:** 10.1101/2023.01.20.524897

**Authors:** Cristina Gil Ávila, Felix S. Bott, Laura Tiemann, Vanessa D. Hohn, Elisabeth S. May, Moritz M. Nickel, Paul Theo Zebhauser, Joachim Gross, Markus Ploner

## Abstract

Biomarker discovery in neurological and psychiatric disorders critically depends on reproducible and transparent methods applied to large-scale datasets. Electroencephalography (EEG) is a promising tool for identifying biomarkers. However, recording, preprocessing, and analysis of EEG data is time-consuming and researcher-dependent. Therefore, we developed DISCOVER-EEG, an open and fully automated pipeline that enables easy and fast preprocessing, analysis, and visualization of resting state EEG data. Data in the Brain Imaging Data Structure (BIDS) standard are automatically preprocessed, and physiologically meaningful features of brain function (including oscillatory power, connectivity, and network characteristics) are extracted and visualized using two open-source and widely used Matlab toolboxes (EEGLAB and FieldTrip). We tested the pipeline in two large, openly available datasets, the LEMON dataset, containing 213 EEG recordings of healthy participants, and the TDBRAIN dataset, containing 1274 EEG recordings, mainly from patients with a psychiatric condition. Additionally, we performed an exploratory analysis of the LEMON dataset that could inspire biomarkers of healthy aging. Thus, the DISCOVER-EEG pipeline facilitates the aggregation, reuse, and analysis of large EEG datasets, promoting open and reproducible research on brain function.

## INTRODUCTION

Biomarkers that relate brain function to cognitive and clinical phenotypes can help in the prediction, treatment, monitoring, and diagnosis of neurological and psychiatric disorders^1, 2^. The successful identification of biomarkers crucially depends on the application of reproducible and transparent methods^3^ to large-scale datasets^4^. Furthermore, to translate biomarkers into clinical practice, they need to be generalizable, interpretable, and easy to deploy in clinical settings.

Electroencephalography (EEG) is a promising tool for biomarker discovery, as it is non-invasive, safe, widely used in clinical and research contexts, portable, and cost-efficient. Consequently, EEG biomarker candidates have been described in depression^5–7^, post-traumatic stress disorder^7, 8^, and chronic pain^9^. Most of these biomarker candidates have been discovered in resting state data, during which spontaneous neural activity is captured. Still, their translation into clinical practice has not yet been successful^1^. This is in part due to small sample sizes and the low availability of objective, transparent, and reproducible EEG preprocessing and analysis methods.

In recent years, notable efforts have been made to automatize, speed up, and increase the transparency of EEG research. A standardized EEG Brain Imaging Data Structure (EEG-BIDS) has been created, which allows for the efficient organization, sharing, and reuse of EEG data^10^. Moreover, automatic preprocessing pipelines have been developed for specific populations, settings, and study designs, including pediatric populations^11, 12^, mobile brain-body imaging^13^, event-related potentials (ERPs)^14, 15^, and generic preprocessing pipelines^15–18^. Beyond, guidelines for the standardized reporting of EEG studies have been established^19^. A logical step is to integrate these solutions into an automatic workflow able to preprocess and extract physiologically informative features in large EEG datasets.

Here, we present DISCOVER-EEG, a comprehensive EEG pipeline for resting state data that extends current preprocessing pipelines by extracting and visualizing physiologically relevant EEG features for biomarker identification. As translation of biomarkers benefits from being neuroscientifically plausible and interpretable^1^, DISCOVER-EEG extracts EEG features, such as oscillatory power, connectivity, and network characteristics, which have been related to brain dysfunction in neurological and psychiatric disorders^20–23^. It builds upon and combines two open-source and widely-used Matlab toolboxes (EEGLAB^24^ and FieldTrip^25^) and adheres to COBIDAS-MEEG guidelines for reproducible MEEG research^19^. It facilitates the aggregation and analysis of large-scale datasets, as it applies to a wide range of EEG setups, and fosters sharing and reusability of the data by handling EEG-BIDS standardized data^10^. Additionally, it can be applied to healthy populations as well as populations with different neuropsychiatric disorders, extending its use to different research and clinical contexts.

We tested DISCOVER-EEG in two large and openly-available datasets, the LEMON dataset^26^, which includes resting state EEG recordings of 213 young and old healthy participants, and the TDBRAIN dataset^27^, which includes resting state EEG recordings of 1274 participants with different psychiatric conditions. We demonstrate the reliability of the pipeline to capture well-known EEG effects, such as the reduction of alpha power during eyes open with respect to eyes closed^28^, in both datasets. Finally, using the LEMON dataset, we present an example analysis investigating differences in EEG features between old and young healthy populations that could inspire biomarkers of healthy aging. Thus, the DISCOVER-EEG pipeline facilitates the preprocessing and analysis of large EEG datasets, promoting open and reproducible research on brain function.

## METHODS

### Design principles

#### Open-source and FAIR code

We developed an automated workflow for fast preprocessing, analysis, and visualization of resting state EEG data (Figure 1) following open science and FAIR principles (Findability, Accessibility, Interoperability, and Reusability)^29^. The code of the DISCOVER-EEG pipeline is published on GitHub and co-deposited at Zenodo, where it is uniquely referenced by a DOI^30^ (*Findability*). The code can be easily downloaded (*Accessibility*) and receive contributions (please refer to the section *Code availability*). To ensure its *Interoperability* and *Reusability*, the pipeline is based on two open-source Matlab toolboxes, EEGLAB^24^ and FieldTrip^25^, which are widely used, maintained, and supported by the developers and the neuroimaging community (i.e., through forums and mailing lists). Basing the pipeline on validated and established software ensures its compatibility with future software updates. Moreover, it facilitates interaction with experts in EEG analysis, who also gave advice and supported the pipeline during its development. The code of the current pipeline is intended to represent a basis that will integrate and benefit from feedback from the neuroimaging community.

**Figure 1.**
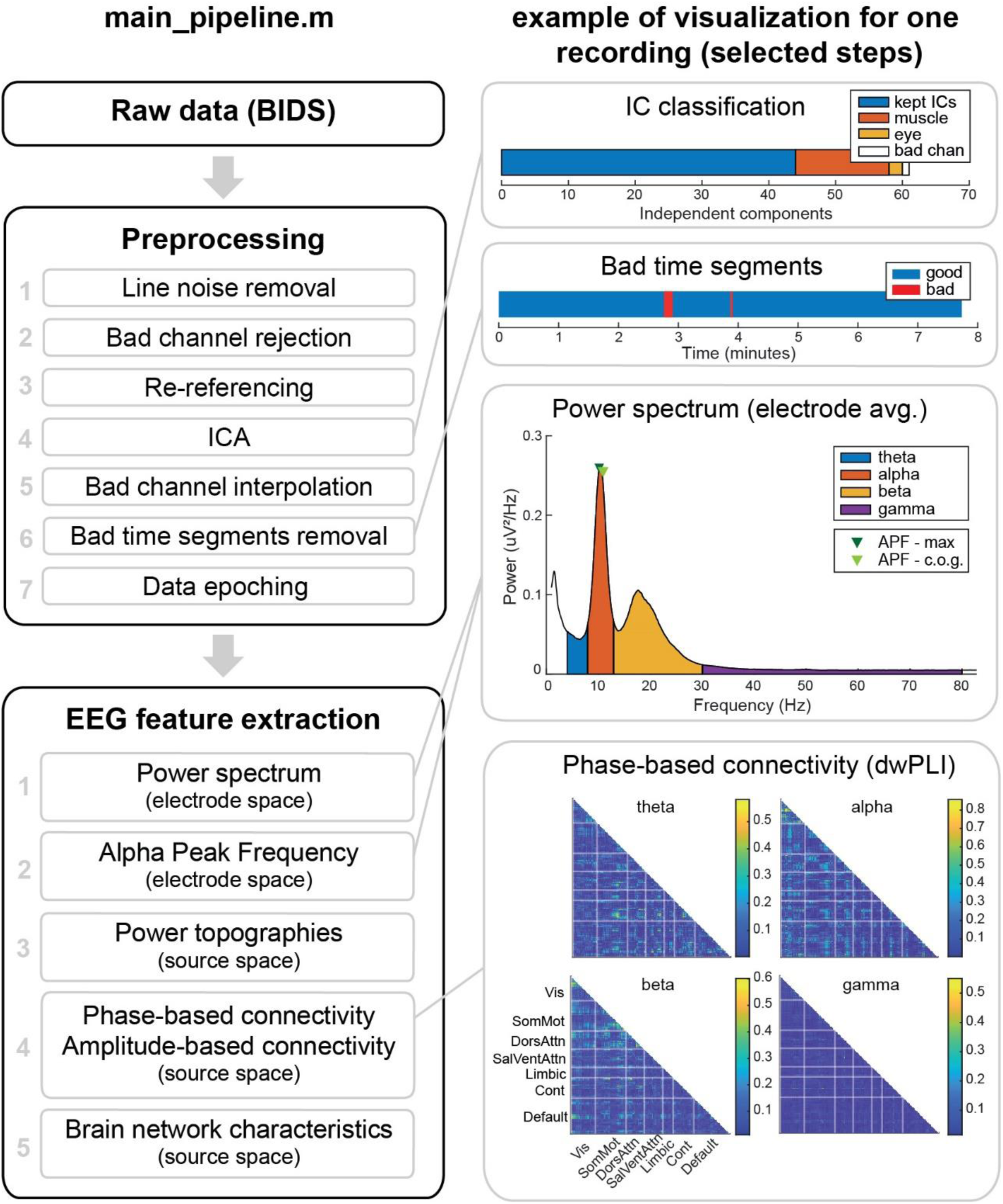
Outline of the DISCOVER pipeline. The left column shows all preprocessing steps and extracted EEG features. The right column shows visualizations for selected steps and features for one EEG recording of the LEMON dataset. APF = Alpha Peak Frequency; BIDS = Brain Image Data Structure; c.o.g. = center of gravity; Cont = Control; DorsAttn = Dorsal Attention; dwPLI = debiased weighted Phase Lag Index; ICA = Independent Component Analysis; SalVentAttn = Salience-Ventral Attention; SomMot = Somato-Motor; Vis = Visual.

#### Data reusability and large-scale data handling

As biomarker discovery needs large datasets, we designed the pipeline in view of data reusability and large-scale data handling. To this end, we followed the FAIR principles of scientific data management^31^ and incorporated the EEG-BIDS standardized data structure^10^ as a mandatory input of the pipeline. Most EEG setups and electrode configurations are compatible with the pipeline thanks to the EEGLAB plugin *bids-matlab-tools.* We refer the reader to the *Results* section for a demonstration of performance across two published datasets with different characteristics, such as participant sample, number of channels, recording length, and sampling rate.

The pipeline was designed for resting state EEG data, during which spontaneous neural activity is recorded. Resting state data can be recorded easily in healthy and patient populations, in different study designs (e.g., cross-sectional, longitudinal), and in different types of neuropsychiatric disorders. Therefore, the use of resting state data facilitates the application to different settings and research questions. EEG recordings might also be accompanied by standardized patient-reported outcomes, such as the PROMIS questionnaires^32^, which can assess symptoms (e.g., pain, fatigue, anxiety, depression) across different neuropsychiatric disorders and, thus, enable cross-disorder analyses. Together, these considerations contribute to the scalability and generalizability of the workflow.

#### Ease of use, transparency, and interpretability

The pipeline consists of a main function, *main_pipeline.m*, in which the preprocessing, feature extraction, and visualization of the data are carried out, and a *params.json* file, in which parameters for preprocessing and feature extraction are defined. This latter file is the only one that needs to be configured by the user and can be easily adapted to dataset and/or user-specific demands. When executing the pipeline, parameters are saved to a separate json file to ensure reproducibility. We recommend DISCOVER-EEG users reporting all parameter configurations as well as software versions when reporting their findings.

Additionally, the pipeline focuses on transparency and interpretability of results. For that reason, single images and an optional PDF report can be generated for each recording to visualize intermediate steps of the preprocessing (Figure 2) and the extracted EEG features (Figure 3). These visualizations can serve as quality control checkpoints and help to detect shorter or corrupted files, misalignment of electrodes, or missing data through fast visual inspection^33^. In that way, we do not exclude any recordings during preprocessing based on quality criteria. Instead, we provide visualization tools that let the users decide how conservative to be in their analysis. Along with their visualization, the preprocessed data and the extracted features of each recording are saved to separate files that can be later used for statistical group analysis and/or biomarker discovery. These output files can be easily imported to other statistical packages or working environments, such as R^34^ or Python^35^, e.g., for applying machine learning or deep learning models.

**Figure 2.**
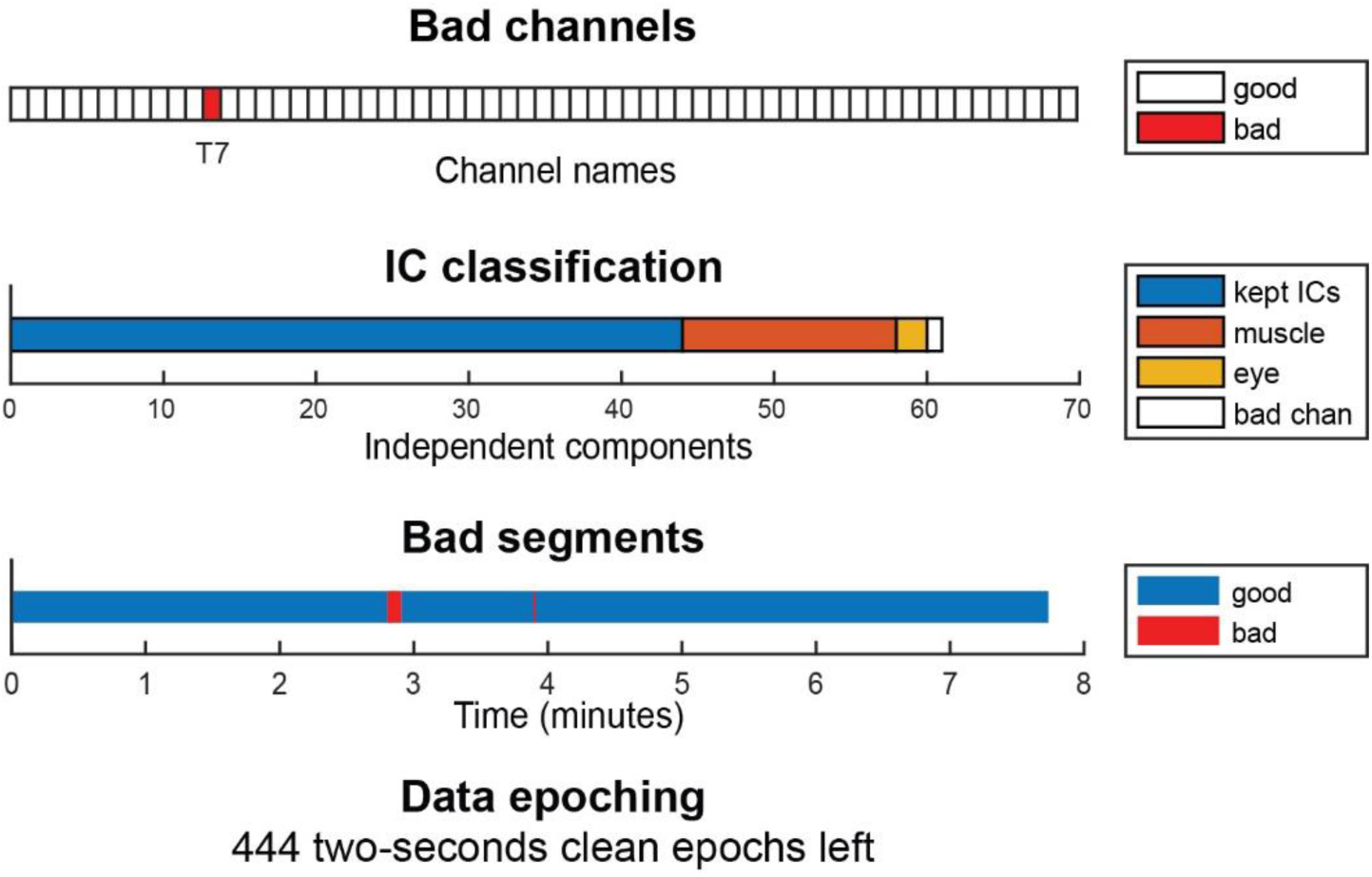
Visualization of the outcome of the preprocessing part of the DISCOVER pipeline. Example of one EEG recording of the LEMON dataset. In the independent component classification (second row), bad channels correspond to the channels that were removed in the bad channel removal step (first row). Bad channels were not included in the independent component analysis. IC = independent component.

**Figure 3.**
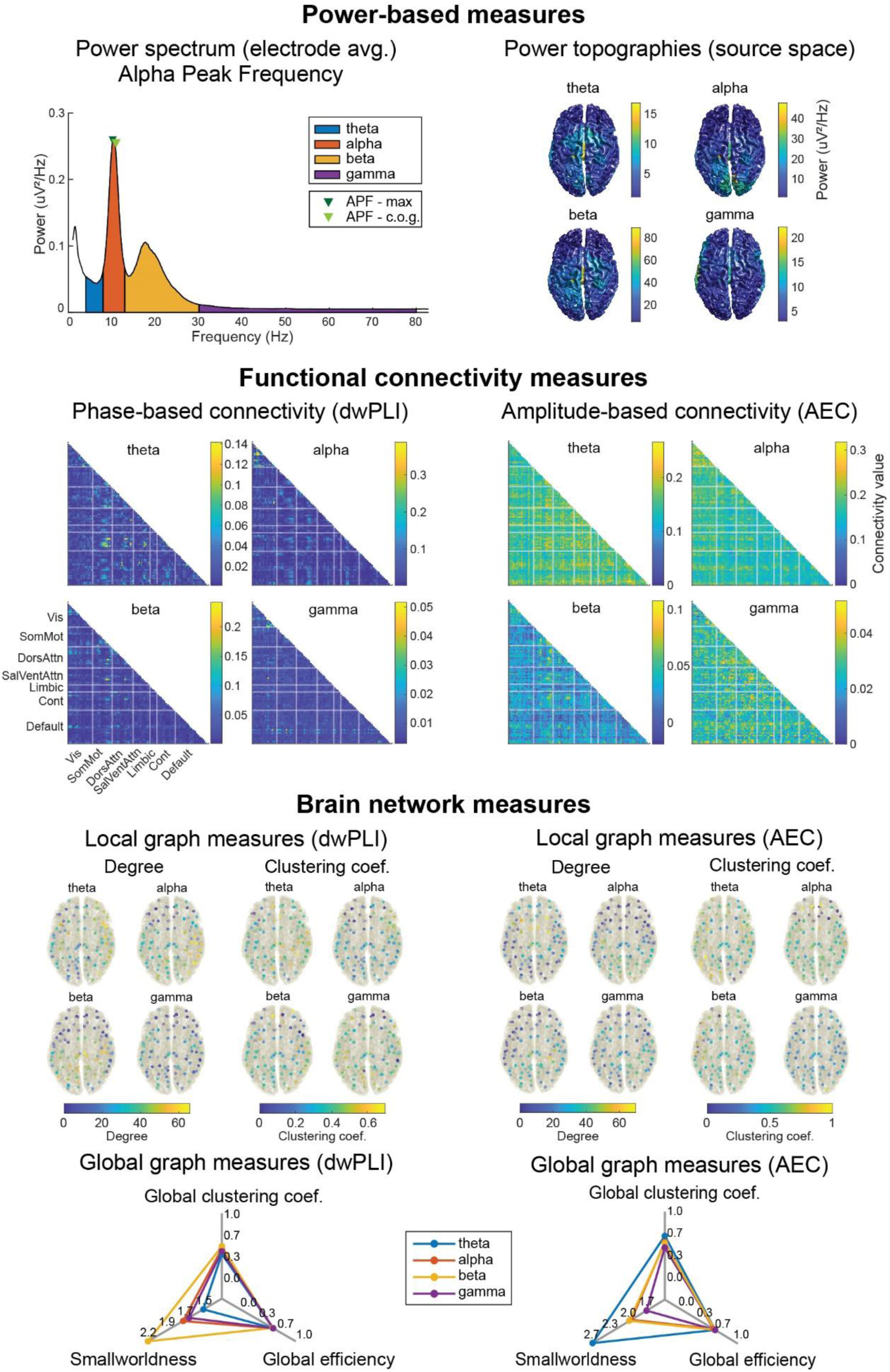
Visualization of the outcome of the feature extraction part of the pipeline. Example for one recording of the LEMON dataset (same recording as Figure 2). **Power-based measures** are extracted in electrode (power spectrum averaged across channels, APF) and source space (power topographies). **Functional connectivity measures** are estimated in source space for 100 pairs of brain regions organized in 7 different functional networks (Visual, Somato-Motor, Salience, Ventral-Attention, Limbic, Control, and Default). Connectivity matrices are symmetric, and thus only lower triangular matrices are shown. **Brain network measures** are characterized by local (degree and clustering coefficient) and global (global clustering coefficient, global efficiency, and smallworldness) graph measures computed on the thresholded connectivity matrices. AEC = Amplitude Envelope Correlation; APF = Alpha Peak Frequency; c.o.g. = center of gravity; Cont = Control; DorsAttn = Dorsal Attention; dwPLI = debiased weighted Phase Lag Index; SalVentAttn = Salience-Ventral Attention; SomMot = Somato-Motor; Vis = Visual.

### Preprocessing

We developed a Matlab-based EEG processing pipeline that complements previous approaches developed in other programming languages, such as the MNE-BIDS-pipeline^18, 36^ in Python. Matlab is widely used by the EEG community, and, in this way, we could use established Matlab-based EEG toolboxes (please refer to the *Design principles* section) which provide robust functions for the computation of functional connectivity measures. Thus, we followed a pragmatic approach towards preprocessing and adopted a simple, established, and automatic workflow in EEGLAB proposed by Pernet et al.^14^ and originally developed for ERP data. We adapted this pipeline to resting-state data and detail the seven preprocessing steps below.

#### Loading the data

The input of the pipeline must be EEG data in BIDS format^10^, including all mandatory sidecar files. We recommend checking the BIDS compliance with the BIDS validator^37^. By default, only EEG channels with standard electrode positions in the 10-5 system^38^ are preprocessed. However, it is possible to preprocess non-standard channels if all electrode positions and their coordinate system are specified in the pertinent BIDS sidecar file. Optionally, after loading, the data can be downsampled to a frequency specified by the user after the application of an anti-aliasing filter.

##### 1. Line noise removal

Line noise is removed with the EEGLAB function *pop_cleanline()*. The CleanLine plugin adaptively estimates and removes sinusoidal artifacts using a frequency-domain (multi-taper) regression technique. CleanLine, as compared to band-stop filters, does not introduce gaps in the power spectrum and avoids the frequency distortions created by filters. This step was added to the Pernet et al.^14^ pipeline to explore brain activity at gamma frequencies (> 30 Hz). The line noise frequency (e.g., 50 Hz or 60 Hz) must be specified in the BIDS file *sub-<label>_eeg.json* to be appropriately removed.

##### 2. High pass filtering and bad channel rejection

Artifactual channels are detected and removed with the function *pop_clean_rawdata().* The first step of this function is the application of a high pass filter with the function *clean_drifts()* with a default transition band of 0.25 to 0.75 Hz. A channel is considered artifactual if it meets any of the following criteria: 1) If it is flat for more than 5 seconds, 2) If the z-scored noise-to-signal ratio of the channel is higher than a threshold set to 4 by default, 3) If the channel time course cannot be predicted from a randomly selected subset of remaining channels at least 80% of the recorded time. Channels marked as artifactual are removed from the data. The mentioned parameters are the defaults proposed by Pernet et al.^14^.

##### 3. Re-referencing

Data is re-referenced to the average reference with the function *pop_reref()*. Optionally, the time course of the original reference channel can be reconstrued and added back to the data if the user specifies it in the file *params.json*. The name of the reference electrode name must be specified in the BIDS file *sub-<label>_eeg.json*.

##### 4. Independent Component Analysis and automatic IC rejection

Independent Component Analysis (ICA) is performed with the function *pop_runica()* using the algorithm *runica*. Artifactual components are automatically classified into seven distinct categories (‘Brain’, ’Muscle’, ’Eye’, ’Heart’, ’Line Noise’, ‘Channel Noise’, and ’Other’) by the ICLabel classifier^39^. Note that ICA is performed only on clean channels, as bad channels detected in step 2 were removed from the data. Therefore, the category ‘Channel Noise’ in the IC classification refers to channel noise remaining after bad channel removal in step 2. By default, only components whose probability of being ‘Muscle’ or ‘Eye’ is higher than 80% are subtracted from the data^14^. Due to the heuristic non-deterministic nature of the ICA algorithm, its results vary across repetitions. That is, every repetition of the ICA algorithm leads to small differences in the reconstructed time series after the removal of artifactual components. While these deviations are small, we noticed that they affect the removal of bad time segments (step 6). For that reason, the pipeline performs by default 10 times steps 4, 5, and 6 and selects the bad time segment mask that is most similar to the average bad time segment mask across all repetitions. This reduces the variability of rejected time segments, but in case of low computational power, the parameter can be set to one repetition in the *params.json* file.

##### 5. Interpolation of removed channels

Channels that were removed in step 2 are interpolated with the function *pop_interp()* using spherical splines^40^. This step was added to the Pernet et al.^14^ pipeline to hold the number of channels constant across participants.

##### 6. Bad time segment removal

Time segments containing artifacts are removed with the Artifact Subspace Reconstruction (ASR) method^41^ implemented in the function *pop_clean_rawdata()*. This method automatically removes segments in which power is abnormally strong. First, a clean segment of data is identified according to the default ASR settings and used for calibration. Calibration data contains all data points in which less than 7.5% of channels are noisy. Here, a channel is defined as noisy if the standard deviation of its RMS is higher than 5.5. Therefore, the length of the calibration data depends on the specific recording. Then, in a sliding window fashion, the whole EEG signal is decomposed via PCA, and the principal subspaces of the window segment are compared with those of the calibration data. Segments with principal subspaces deviating from the calibration data (20 times higher variance) are removed. Again, default parameters are in line with Pernet et al.^14^.

##### 7. Data segmentation into epochs

Lastly, the continuous data are segmented into epochs with the function *pop_epoch().* By default, data are segmented into 2-second epochs with a 50% overlap. Although longer epochs might be desirable for the Alpha Peak Frequency estimation to increase frequency resolution, short epochs favor the reliability of functional connectivity measures^42, 43^. Thus, we propose 2- second epochs to establish a balance between frequency resolution, stationarity of the signal, and reliability of the later extracted features. Fifty percent overlap was chosen to provide a smooth estimation of the power spectra and mitigate the loss of signal due to tapering^44^. Epochs containing a discontinuity (e.g., because a segment containing an artifact was discarded) are rejected automatically. Data segmentation was adapted from Pernet et al.^14^, which focused on event-related data.

#### EEG feature extraction

DISCOVER-EEG extracts EEG features that have been related to different neurological and psychiatric disorders and have the potential to turn them into neuroscientifically plausible and interpretable biomarkers.

In electrode space, power spectra and the Alpha Peak Frequency, i.e., the frequency at which a peak in the power spectrum in the alpha range occurs, are extracted. Power changes in different frequency bands have been found in a broad spectrum of neuropsychiatric disorders^20^, and APF has been correlated with behavioral and cognitive characteristics^45^ in aging and disease^46^.

In source space, two measures of functional connectivity at the theta, alpha, beta, and gamma bands, are extracted. Additionally, brain networks derived from these connectivity matrices are further characterized with common graph theory measures^47^. Functional connectivity is a promising biomarker candidate, as it has been successfully used for the classification, tracking, and stratification of patients with several neurological and psychiatric disorders, such as Alzheimer’s disease^48^, Post Traumatic Stress Disorder^8^, and Major Depressive Disorder^7, 49^ (for a recent review see^50^). In EEG, functional connectivity measures are commonly classified as either phase-based or amplitude-based, each type capturing different and complementary communication processes in the brain^51^. This pipeline thus includes a phase-based measure, the debiased weighted Phase Lag Index (dwPLI)^52^, and an amplitude-based measure, the orthogonalized Amplitude Envelope Correlation (AEC)^53^. Both functional connectivity measures are undirected and have low susceptibility to volume conduction. The dwPLI also has the advantage of being more sensitive and capable of capturing non-linear relationships compared to other phase-based measures^44^. For each connectivity matrix, two local graph measures (the degree and the clustering coefficient) are calculated at each source location, and three global graph measures (the global clustering coefficient, the global efficiency, and the smallworldness) summarize the whole network in one value.

Power and connectivity features are computed using the preprocessed and segmented data in FieldTrip. Graph theory measures are computed with the Brain Connectivity Toolbox^47^.

Specific parameters on feature extraction are defined in the file *params.json* and detailed below.

##### 1. Power spectrum

Power spectra are computed with the FieldTrip function *ft_freqanalysis* between 1 and 100 Hz using Slepian multitapers with +/-1 Hz frequency smoothing. For 2-second epochs, the maximum frequency resolution of the power spectrum is, by definition,n the inverse of the epoch length, I,.e. 0.5 Hz. As frequency band limits are determined by the spectral resolution, the pipeline zero-pads the epochs to 5 seconds, which yields a resolution of 0.1 Hz, to better capture the frequency range of the bands. Thus, the frequency band limits for theta, alpha, beta, and gamma are 4 to 7.9 Hz, 8 to 12.9 Hz, 13 to 30 Hz, and 30.1 to 80 Hz, respectively. These limits are defined by the COBIDAS-MEEG^19^ guidelines and maintained throughout all features. Power spectra are computed for every channel and then averaged across epochs and channels to obtain a single global power spectrum. This global power spectrum is saved to *sub-<label>_power.mat* files and visualized with the different frequency bands highlighted in different colors (Figure 3).

##### 2. Alpha Peak Frequency

APF is computed based on the global power spectrum in the alpha range (8 to 12.9 Hz). There are two widely accepted strategies to assess the APF in the literature^54^. The first one reports the frequency at which the highest peak in the alpha range occurs (peak maximum). This comes with the problem that not always a peak is present in the power spectrum. Therefore, the second strategy, calculating the center of gravity (c.o.g.) of the power spectra in the alpha band, is also frequently used^55^. DISCOVER-EEG reports both measures for estimating APF. The peak maximum is computed with the Matlab function *findpeaks()*. If there is no peak in the alpha range, no value is returned. The center of gravity is computed as the weighted average of frequencies in the alpha band, each frequency being weighted by their power^55^. Both measures are visualized in the same figure as the global power spectrum (Figure 3). Individual APF values are also saved to the *sub-<label>_peakfrequency.mat* files.

##### 3. Source reconstruction

To mitigate the volume conduction problem when computing functional connectivity measures^56^, we perform a source reconstruction of the preprocessed data with an atlas-based beamforming approach^57^. For each frequency band, the band-pass filtered data from sensor space is projected into source space using an array-gain Linear Constrained Minimum Variance (LCMV) beamformer^58^. As source model, we selected the centroids of 100 regions of interest (ROIs) of the 7-network version of the Schaefer atlas^59^. This atlas is a refined version of the Yeo atlas and follows a data-driven approach in which 100 parcellations are clustered and assigned to 7 brain networks (Visual, Somato-Motor, Dorsal attention, Salience-Ventral attention, Limbic, Control, and Default networks). This atlas can be easily changed according to the user’s preferences in the *params.json* file. The lead field is built using a realistically shaped volume conduction model based on the template of the Montreal Neurological Institute (MNI) available in FieldTrip (*standard_bem.mat*) and the source model. Spatial filters are finally constructed with the covariance matrices of the band-passed filtered data and the described lead fields. A 5% regularization parameter is set to account for rank deficiencies in the covariance matrix, and the dipole orientation is fixed to the direction of the maximum variance following the most recent recommendations^60^. An estimation of the power for each source location is obtained using the spatial filter and band-passed data. A visualization of the source power in each frequency band (Figure 3) is provided by projecting the band-specific source power to a cortical surface model provided as a template in FieldTrip (*surface_white_both.mat*).

##### 4. Functional connectivity

After the creation of spatial filters in the four frequency bands, virtual time series in the 100 source locations are reconstructed for each frequency band by applying the respective band-specific spatial filter to the band-pass filtered sensor data. Then, the two functional connectivity measures (dwPLI and AEC) are computed for each frequency band and for each combination of the 100 reconstructed virtual time series. Average connectivity matrices for each band are visualized (Figure 3) and saved to separate files (*sub-<label>_<conmeasure>_<band>.mat*). The phase-based connectivity measure dwPLI is computed using the FieldTrip function *ft_connectivityanalysis* with the method *wpli_debiased*, which requires a frequency structure as input. Therefore, Fourier decompositions of the virtual time series are calculated in each frequency band with a frequency resolution of 0.5 Hz. Thereby, a connectivity matrix is obtained for each frequency of interest in the current frequency band. Connectivity matrices are then averaged across each frequency band resulting in one 100×100 connectivity matrix for each of the four frequency bands.

The amplitude-based connectivity measure AEC is computed according to the original equations with a custom function *compute_aec*, as the original implementation was not available in FieldTrip. For each epoch, the analytical signal of the virtual time series is extracted at each source location with the Hilbert transform. For each source pair, the analytical signal at source A is orthogonalized with respect to the analytical signal at source B, yielding the signal A_⟂B_. Then, the Pearson correlation is computed between the amplitude envelope of signals B and A_⟂B_. To obtain the average connectivity between sources A and B, the Pearson correlation between the amplitude envelopes of analytical signal A and signal B_⟂A_ is also computed, and the two correlation coefficients are averaged. In this way, we obtain a 100×100 connectivity matrix for each epoch. We finally average the connectivity matrices across epochs, resulting in one 100×100 connectivity matrix for each of the four frequency bands.

##### 5. Brain network characteristics

Graph measures were computed on thresholded and binarized connectivity matrices^47^. Matrices were binarized by keeping the 20% strongest connections, as this threshold delivers fairly reproducible graph measures based on dwPLI and AEC connectivity^61^. Nevertheless, it is good practice to test the reliability of final results with different binarizing thresholds^62^. This threshold can be easily changed in the file *params.json*.

The computed local network measures are the degree and the clustering coefficient. The degree is the number of connections of a node in the network. The clustering coefficient is the percentage of triangle connections surrounding a node. The measures, thus, assess the global and local connectedness of a node, respectively.

The computed global network measures are the global clustering coefficient, the global efficiency, and the smallworldness. The global clustering coefficient is a measure of functional segregation in the network and is defined as the average clustering coefficient of all nodes. Global efficiency is a measure of functional integration in the network and is defined as the average of the inverse shortest path length between all pairs of nodes. High global efficiency indicates that information can travel efficiently between regions that are far away. Smallworldness compares the ratio between functional integration and segregation in the network against a random network of the same size and degree. Smallworld networks are highly clustered and have short characteristic path length (average shortest path length between all nodes) compared to random networks^63^. Local and global measures are visualized per recording and saved to files named *sub-<label>_graph_<conmeasure>_<band>.mat* (Figure 3).

## RESULTS

### Testing DISCOVER-EEG in two large, public datasets

We tested the DISCOVER-EEG pipeline in two well-documented and openly-available resting state EEG datasets, the LEMON dataset^26, 64^, including 213 healthy participants, and the TD- BRAIN dataset^27, 65^, including 1274 participants, most of them with different psychiatric conditions. Below we detail the characteristics of both datasets and the outcome of their preprocessing with the DISCOVER-EEG pipeline during the eyes closed and eyes open resting state. These results demonstrate that the pipeline is robust across different EEG systems, electrode numbers and layouts, sampling rates, and recording lengths.

We additionally show that the pipeline can capture a well-known neurophysiological phenomenon in both datasets, the difference in oscillatory alpha power between eyes open and eyes closed^28^. A comparison of power spectra between eyes open and eyes closed had previously been reported for the TD-BRAIN dataset to prove the neurophysiological validity of the data^27^, but not for the LEMON dataset. Reproducing this effect with a different preprocessing strategy for the TD-BRAIN dataset and confirming it for the first time for the LEMON dataset, proves the reliability of the DISCOVER-EEG pipeline to capture well-known features of EEG data.

#### LEMON dataset

The LEMON dataset is a publicly available resting state EEG dataset of 213 young and old healthy participants acquired in Leipzig (Germany) to study mind-body-emotion interactions^26^. This dataset is divided into two groups based on the age of the participants: a ‘young’ group (20 to 35 years old; N = 143; 43 females) and an ‘old’ group (55 to 80 years old; N = 70; 35 females). One participant (male) had an intermediate age and was not included in the posterior statistical analyses (next Results section). Resting state EEG was recorded with a BrainAmp MR plus amplifier using 62 active ActiCAP electrodes (61 scalp electrodes in the 10-10 system positions and 1 VEOG below the right eye) provided by Brain Products Gmbh, Gilching, Germany. The ground electrode was located at the sternum, and the reference electrode was FCz. The recording sampling rate was 2500 Hz. Recordings contained 16 blocks of 1-minute duration, 8 with eyes closed and 8 with eyes open in an interleaved fashion. Before the execution of the pipeline, all blocks corresponding to eyes closed and eyes open conditions were extracted and concatenated by condition, including boundaries between blocks. In both conditions, two recordings were discarded because the data was truncated. In the eyes closed condition, one recording was discarded because there were no eyes closed marker events. Therefore, the final sample sizes were 210 recordings for eyes closed and 211 for eyes open. For both conditions, recordings of 8 minutes duration were entered into the pipeline.

#### TD-BRAIN dataset

The TD-BRAIN dataset is a publicly available, heterogenous resting state EEG dataset aggregated to obtain neurophysiological insights in psychiatric disorders. It includes 1274 participants, most of them suffering from a psychiatric disorder. The primary diagnoses are Major Depressive Disorder (N = 426), Attention Deficit Hyperactivity Disorder (N=271), Subjective Memory Complains (N=119), and obsessive-compulsive disorder (N=75). The dataset also includes healthy participants (N=47) and participants with unknown diagnoses (N=255). Resting state EEG was recorded with a Compumedics Quickcap or ANT-Neuro Wavegurard Cap using 26 EEG Ag/AgCl electrodes positioned in the 10-10 system. Additionally, five electrodes recorded vertical and horizontal eye movements, one electrode recorded muscle activity at the masseter, and one electrode captured the electrocardiogram at the clavicle bone. Ground electrode was placed at AFz, and recordings were referenced offline to averaged mastoids (A1 and A2). Recording sampling was 500 Hz. Resting state was acquired with eyes closed and eyes open in two respective blocks of 2 minutes duration. Some participants had more than one recording session, but only one recording per participant was included in the pipeline. The final sample sizes were 1274 recordings, both for eyes closed and open.

#### DISCOVER-EEG preprocessing

Table 1 and Figure 4 show an overview of the DISCOVER-EEG preprocessing results, comparing datasets with different numbers of EEG electrodes (61 for the LEMON and 26 for the TD-BRAIN dataset) and recording lengths (8 minutes for the LEMON and 2 minutes for the TD-BRAIN datasets). These results point to a fair amount of data remaining after preprocessing and are in the same range as other automatic preprocessing pipelines^11, 16^. They could be useful in future studies interested in benchmarking different EEG configurations or EEG preprocessing strategies.

**Figure 4.**
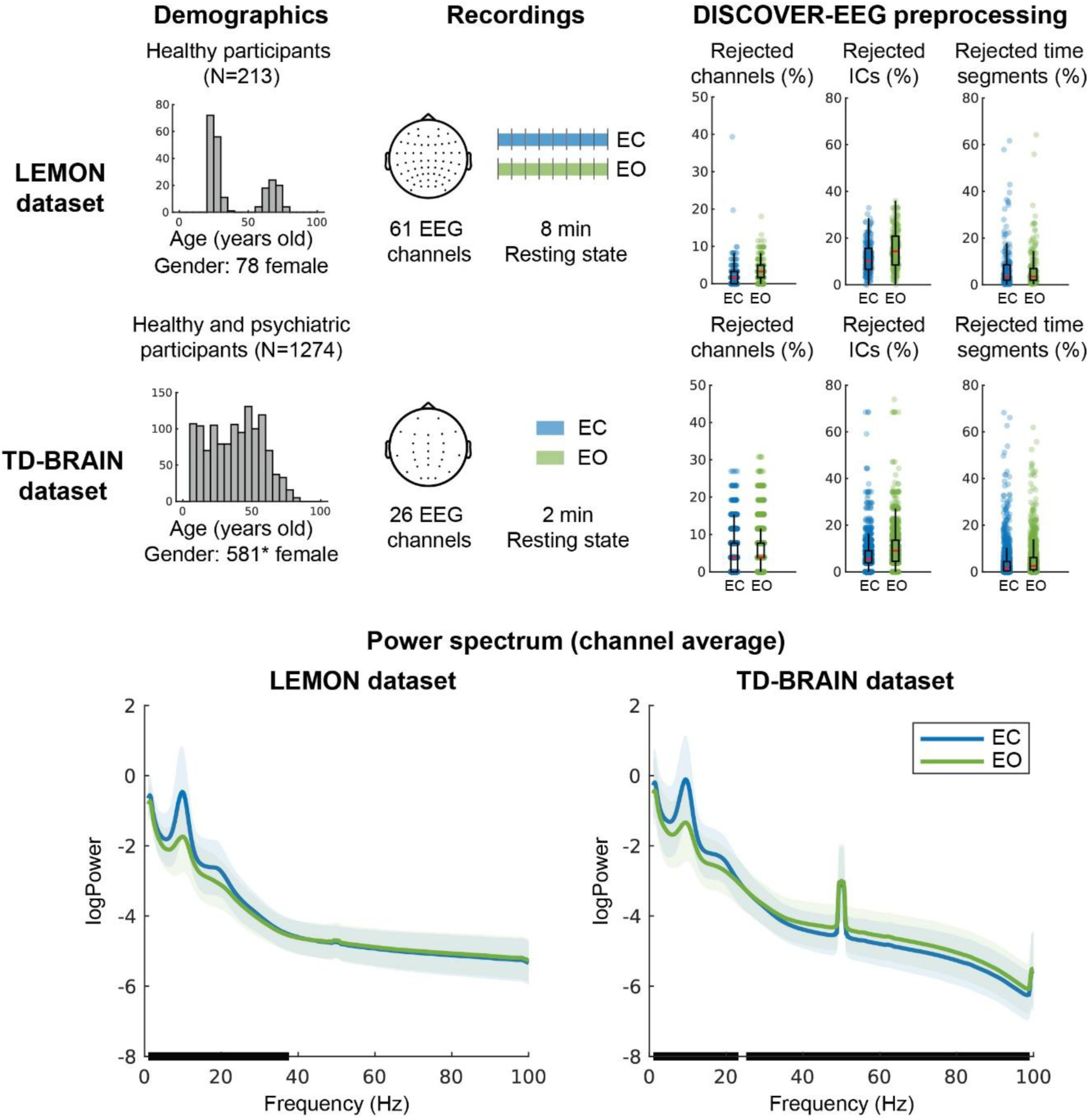
Overview of the LEMON and TD-BRAIN datasets, results of their preprocessing for resting state eyes closed (blue) and open (green), and average power spectra across participants and channels. **Demographics** includes a histogram depicting the age of the participants in bins of 5 years. **Recordings** depicts the EEG setup layout and the recordings’ average duration. Vertical bars in the recording duration of the LEMON dataset represent the boundaries between the concatenated 1-minute blocks in which the data was acquired. **DISCOVER-EEG preprocessing** presents the preprocessing variables expressed in percentage relative to the number of electrodes (rejected channels, rejected ICs) or length of the recording (rejected time segments). Boxplots visualize the distribution of these variables. The median is indicated by a red horizontal line, and the first and third quartiles are indicated with black boxes. The whiskers extend to 1.5 times the interquartile range. Blue dots overlaid to the boxplots represent the individual recordings of each dataset. **Power spectrum** depicts the grand average power spectra across participants and channels for eyes open and eyes closed conditions. Shaded areas indicate the standard deviation of the power spectra across participants. Thick black lines over the x axis indicate significant differences between conditions (dependent samples cluster-based permutation test). Note that in the TD-BRAIN dataset, line noise could not be completely removed due to high levels of noise in some recordings. EC = Eyes Closed. EO = Eyes Open. ICs = Independent Components. *The gender of 18 participants was missing in the TD-BRAIN dataset.

**Table 1.**
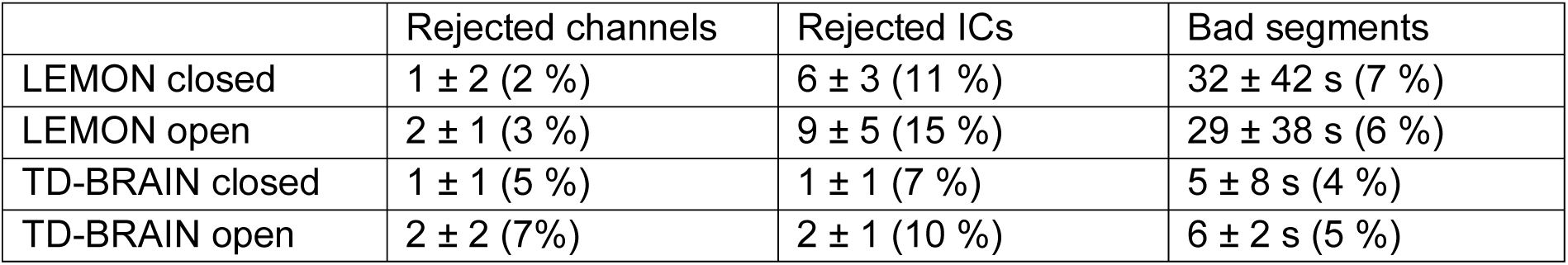
DISCOVER-EEG preprocessing summary of the LEMON and TD-BRAIN datasets for eyes open and closed conditions. Average ± standard deviation across participants is stated for the number of rejected channels, number of rejected ICs, and length of rejected bad time segments in seconds. The average of each variable in percentage is stated between brackets. The maximum number of ICs varied between participants as it depended on the number of channels that were rejected for each participant. ICs = Independent components.

|  | Rejected channels | Rejected ICs | Bad segments |
| --- | --- | --- | --- |
| LEMON closed | 1 ± 2 (2 %) | 6 ± 3 (11 %) | 32 ± 42 s (7 %) |
| LEMON open | 2 ± 1 (3 %) | 9 ± 5 (15 %) | 29 ± 38 s (6 %) |
| TD-BRAIN closed | 1 ± 1 (5 %) | 1 ± 1 (7 %) | 5 ± 8 s (4 %) |
| TD-BRAIN open | 2 ± 2 (7%) | 2 ± 1 (10 %) | 6 ± 2 s (5 %) |

#### DISCOVER-EEG validation

As proof of validation, we replicated the well-known physiological effect of alpha power attenuation in eyes open resting state with respect to eyes closed in both datasets. We estimated the power spectrum for each recording and condition as described in the feature extraction section. For visualization purposes, we performed a grand average across participants and channels for each dataset and condition (Figure 4). To test for differences between conditions, we performed a dependent samples cluster-based permutation test across frequencies in the range of 1 to 100 Hz. The tests were two-tailed, and the cluster threshold was set to 0.05. In the LEMON dataset, a significant positive cluster, indicating higher power values in the eyes closed condition, was found in the 1 to 37.2 Hz range (p = 0.002, cluster statistic = 1310). In the TD-BRAIN, a positive cluster was found in the 1 to 23.2 Hz range (p = 0.002, cluster statistic = 2037), and a negative cluster in the 25.2 to 99 Hz range (p = 0.002, cluster statistic = -3254). These results indicate lower power in the eyes open condition in the alpha range (8 to 12.9 Hz) for both datasets.

### Example analysis with DISCOVER-EEG

To exemplify how DISCOVER-EEG features could be aggregated and analyzed, we conducted an exploratory analysis in the LEMON dataset to investigate age-related differences in resting state EEG. Recently, the measure of brain age, i.e., the expected level of cognitive function of a person with the same chronological age, has been proven a valuable marker of cognitive decline in healthy and clinical populations^66, 67^. Brain age is usually estimated by machine learning models trained to predict the chronological age of a participant based on neuroimaging data. One criticism of these methods is their limited neuroscientific interpretability, as it is not straightforward which features the models used to predict brain age^66^. It is, therefore, highly useful to explore which physiologically meaningful brain features change with age.

To this end, we statistically compared the features automatically extracted by the pipeline in the ‘old’ and ‘young’ groups using Bayesian statistics performed in Matlab with the bayesFactor package^68^ (Figure 5). We also provide the scripts used to perform this group analysis and the visualization of group-level data together with the pipeline code.

**Figure 5.**
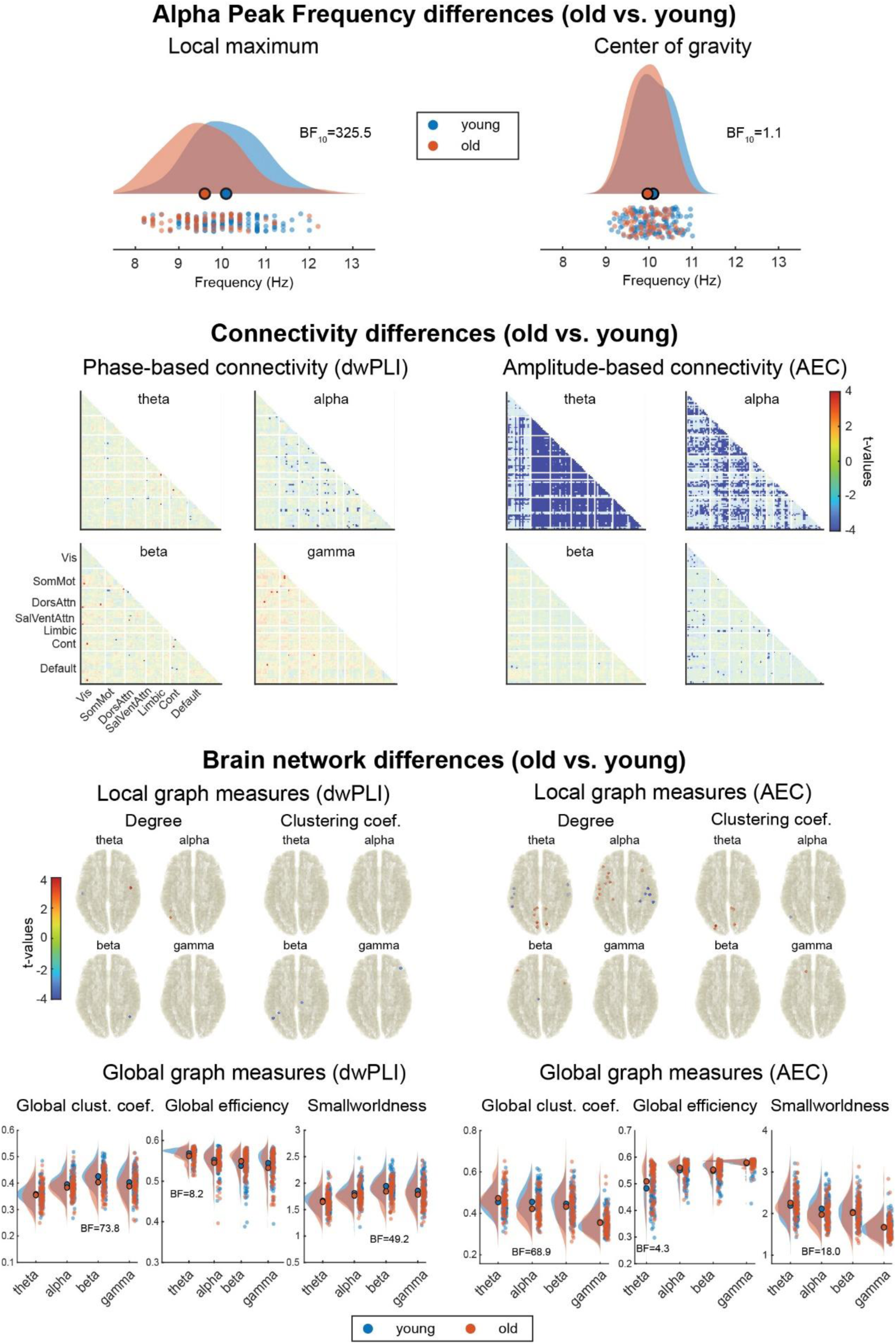
Age-related differences in resting state EEG between old and young participants of the LEMON dataset. **Alpha Peak Frequency differences** between the old and the young group are visualized using raincloud plots^70^. **Connectivity differences** between the old and young groups. Blue values indicate lower connectivity in the old group. All connections not showing strong evidence for or against a connectivity difference (1/30 < BF_10_ < 30) are faded out. **Brain network differences** between the old and young group. With regard to local graph measures, only locations with strong evidence for (BF_10_ > 30) or against (BF_10_ < 1/30) a difference in the graph measures are depicted. Only results in favor of a difference between groups were found (blue dots indicate lower local graph measures in the old group, and red dots indicate higher local graph measures in the old group). With regard to global graph measures, only BF_10_ showing substantial evidence for (BF_10_ > 3) or against (BF_10_ < 1/3) the alternative hypothesis were included as insets to facilitate the reading. AEC = Amplitude Envelope Correlation; BF = Bayes Factor; Cont = Control; DorsAttn = Dorsal Attention; dwPLI = debiased weighted Phase Lag Index; SalVentAttn = Salience-Ventral Attention; SomMot = Somato-Motor; Vis = Visual

We tested whether old participants had different APF values than young participants with two-sided independent samples Bayesian t-tests. Two tests were performed, one for each APF measure (the local maximum and the center of gravity). Results show very strong evidence in favor of the alternative hypothesis, i.e., a lower APF in the old compared to the young group, when the APF is computed as local maximum peak (BF_10_ = 325.5), but inconclusive evidence when the APF is computed as the center of gravity (BF_10_ = 1.1) (Figure 5, first row).

We further tested whether there were differences in the connectivity matrices between young and old participants. For each connectivity measure (dwPLI and AEC) and frequency band (theta, alpha, beta, and gamma), we compared the connectivity values of each undirected source pair between the young and the old groups. Thus, we performed 9900 two-sided independent samples Bayesian t-tests per connectivity matrix. In Figure 5, second row, we depict t-values color-coded to show the direction of effects. Statistical tests showing strong evidence in favor (BF_10_ > 30) or against (BF_10_ < 1/30) of the alternative hypothesis are not faded out. We observed strong evidence in favor of a reduction of phase-based connectivity in old participants, predominantly in the alpha band as well as a reduction of amplitude-based connectivity (Figure 5, second row, non-masked blue values).

We finally tested whether there were any differences between old and young participants in the graph measures. For the local graph measures, we performed one test per source location, i.e., 100 independent sample Bayesian t-tests for each graph measure, connectivity measure, and frequency band. For the global measures, we performed a two-sided independent sample Bayesian t-test per graph measure, connectivity measure, and frequency band. With regard to global measures, the most prevalent differences appeared at low frequencies (theta and alpha) using the AEC (Figure 5, third row, blue and red dots in brain sketches have BF_10_ > 30). The strongest effects with regard to global measures are a reduction of global efficiency and smallworldness of the older group in the beta band for the dwPLI and the alpha band for the AEC (Figure 5, third row, raincloud plots with inset BF_10_ indicating strong evidence). Together, these results show a reduction of local connectivity at theta and alpha frequencies and an increase in network integration at alpha and beta frequencies in older participants.

Overall, the current findings are in line with previous EEG literature, which has reported a slowing of APF and a general decrease in functional connectivity and network integrity in older individuals^46, 69^.

This example analysis could inspire future studies aimed at discovering explainable biomarkers of healthy aging or risk biomarkers of cognitive decline. However, it does not intend to present a ready-to-use biomarker for aging, which would require validation beyond the scope of this manuscript.

## DISCUSSION

Here, we present DISCOVER-EEG, an open and fully automated pipeline that enables fast and easy aggregation, preprocessing, analysis, and visualization of resting state EEG data. The current pipeline builds upon state-of-the-art automated preprocessing elements and extends them by including the computation and visualization of physiologically relevant EEG features. These EEG features follow recent COBIDAS guidelines for MEEG research^19^, are implemented in widely used EEG toolboxes, and have been repeatedly associated with cognitive and behavioral measures in healthy as well in neuropsychiatric populations. Therefore, they could represent promising biomarker candidates for neurological and psychiatric disorders.

Importantly, this pipeline presents one out of many ways of preprocessing and analyzing resting state data and is not intended to be the ultimate solution for EEG analysis. Instead, it aims to represent a reasonable and pragmatic realization for accelerating the acquisition, preprocessing, and analysis of large-scale datasets with the potential to discover physiologically plausible and interpretable biomarkers. To this end, we adopted a previously published, simple, and robust preprocessing strategy and selected a well-defined set of EEG features that could be useful for biomarker discovery. However, this pipeline might not suit all populations, settings, or research paradigms, such as children, real-life ecological EEG assessments, or paradigms assessing event-related potentials. Additionally, specific EEG layouts, such as those with few electrodes and limited scalp coverage, might not be adequate for source localization or average referencing. However, it is compatible with different types of EEG systems, and its focus on resting state data makes it suitable for patients and healthy populations. The features extracted by DISCOVER-EEG represent a basis for developing neurophysiologically plausible and interpretable biomarkers of neurological and psychiatric disorders. They can be easily extended and adapted to specific study designs, as the modular configuration of the code allows for substituting, removing, or adding specific steps of the preprocessing and feature extraction.

By default, the pipeline uses a realistically shaped volume conduction model based on the template of the Montreal Neurological Institute (MNI) for source localization. For optimal accuracy of source reconstruction, individual MRIs or at least individual electrode positions could be recorded along with the EEG data. However, this would substantially increase the effort and time needed to acquire data and would hinder the fast generation of large new datasets. For that reason, the pipeline uses a generic template for source localization. However, it might be desirable to have different templates that better reflect the variability of head shapes in the future.

On a broader scope, it should always be considered whether datasets used for biomarker discovery are representative of the population of interest or whether they are biased towards young, Caucasian, highly educated populations, as is often the case^71^. The participants of the studies used here to test the pipeline were recruited based on convenience sampling and, therefore, might not cover the entire population. The creation and adoption of data standards such as BIDS will help to mitigate this fact by promoting collaboration and data sharing around the world.

Our intention with this pipeline was to push the field of EEG biomarker discovery forward to the acquisition and analysis of large datasets, as needed in neuroimaging and artificial intelligence. Moreover, the provided example analysis can serve as a starting point for researchers who want to use complex measures of brain function. However, we do not claim to present a ready-to-use biomarker for aging, nor that the approach we propose is the best or unique way to develop such a biomarker. Instead, it is intended to represent an offer to the community to assess physiologically plausible and interpretable EEG features in an efficient, transparent, and reproducible manner. Therefore, it can help the discovery of EEG-based biomarkers in neuropsychiatric disorders and promotes and facilitates open and reproducible assessments of brain function in EEG communities and beyond.

## DATA AVAILABILITY

The LEMON dataset^64^ was accessed via http from the Max Plax Institute Leipzig webpage, and it is described in an accompanying publication^26^.

The TDBRAIN dataset^65^ was accessed from the webpage of Brainclinics Foundation, and it is described in an accompanying publication^27^.

## CODE AVAILABILITY

The EEG pipeline code is available at GitHub under the CC-BY 4.0 license, and it is co-deposited in Zenodo, and referenced with a unique DOI^30^.

The pipeline was created and tested in Matlab 2020b (The Mathworks, Inc.) on Ubuntu 18.04.5 LTS with the Signal Processing and Statistical and Machine Learning Toolboxes installed. EEGLab (v2022.0)^24^ with the plugins bids-matlab-tools (v6.1), bva-io (v1.7), firfilt, (v2.4), cleanLine (v2.0), ICLabel (v1.3), clean_rawdata (v2.6) and dipfilt (v4.3) were installed and used for preprocessing. FieldTrip (revision ee916f5e5)^25^ was used for source reconstruction and EEG feature extraction, and the Brain Connectivity Toolbox (version 03 2019)^47^ was used for network analysis.

## ACKNOWLEDGMENTS

The study has been supported by the TUM Innovation Network Neurotechnology for Mental Health (NEUROTECH), the Deutsche Forschungsgemeinschaft (PL321/14-1) and the TUM School of Medicine (KKF).

## AUTHOR CONTRIBUTIONS

`-` Conceptualization: C.G.A. and M.P.

`-` Methodology: C.G.A. and M.P.

`-` Software: C.G.A., F.S.B., J.G.

`-` Validation: C.G.A., F.S.B., J.G.

`-` Formal analysis: C.G.A.

`-` Investigation: C.G.A. and P.T.Z.

`-` Data curation: C.G.A.

`-` Visualization: C.G.A. and M.P.

`-` Writing – Original draft: C.G.A. and M.P.

`-` Writing – Review and editing: C.G.A., F.S.B., L.T., V.D.H., E.S.M., M.M.N., P.T.Z., J.G. and M.P.

`-` Supervision: M.P.

`-` Project administration: M.P.

- Funding acquisition: M.P

